# Revisiting the *Ancylostoma caninum* secretome provides new information on hookworm-host interactions

**DOI:** 10.1101/139006

**Authors:** Taylor Morante, Catherine Shepherd, Constantin Constantinoiu, Alex Loukas, Javier Sotillo

## Abstract

Hookworm infection is a major tropical parasitic disease affecting almost 500 million people worldwide. These soil-transmitted helminths can survive for many years in the intestine of the host, where they feed on blood, causing iron deficiency anaemia and other complications. To avoid the host’s immune response the parasite releases excretory/secretory products (ESPs), a complex mixture of glycans, lipids and proteins that represent the major host-parasite interface. Using a combination of separation techniques such as SDS-PAGE and OFFGEL electrophoresis, in combination with state-of-the-art mass spectrometry we have reanalysed the dog hookworm, *Ancylostoma caninum*, ESPs (*Ac*ES). We identified 315 proteins present in the *Ac*ES, compared with just 105 identified in previous studies. The most highly represented family of proteins is the SCP/TAPs (90 of the 315 proteins), and the most abundant constituents of *Ac*ES are homologues of the tissue inhibitors of metalloproteases (TIMP) family. We identified putative vaccine candidates and proteins that could have immunomodulatory effects for treating inflammatory diseases. This study provides novel information about the proteins involved in host-hookworm interactions, and constitutes a comprehensive dataset for the development of vaccines and the discovery of new immunoregulatory biologics.

---

Soil-transmitted helminthiases (including trichuriasis, ascariasis and hookworm infections) are debilitating parasitic diseases that affect more than two billion people worldwide [1], with increased incidence occurring in impoverished and underdeveloped societies. Hookworms alone affect almost 500 million people in tropical regions of South America, Africa and Asia [2], and chronic infections result in iron-deficiency anaemia and even physical and intellectual retardation in young children [3]. Adult hookworms live in the intestine of vertebrate hosts where they feed on blood, and constantly release products into their surrounding environment through excretion and secretion mechanisms (excretory/secretory products, ESPs). The ESPs contain proteins that facilitate a parasitic existence, notably penetration of and migration within a host, feeding on host tissues, and evasion of the host immune response [4]. In addition, recently, hookworm ESPs have been shown to contain immunoregulatory properties that can protect mice against inflammatory diseases such as inflammatory bowel diseases and asthma [5–9].

Due to the difficulty in obtaining samples from the human hookworm *Necator americanus*, the dog hookworm *Ancylostoma caninum* has been extensively used as a model to study hookworm infections. The first proteomic characterisation of the ESPs produced by *A. caninum* (*Ac*ESP) was performed by Mulvenna et al. in 2009 [10]; however, herein we revisit this data since the *A. caninum* genome was not available at the time and the sensitivity of mass spectrometers has improved dramatically since this last study was conducted.

A total of ∼300 *A. caninum* adult worms were obtained from the small intestine of 5 fresh cadaver dogs that had been naturally infected. Worms were divided into two different batches and incubated in 5x substrate (Dulbecco’s Phosphate Buffered Saline (DPBS) (+) CaCl_2_ (+) MgCl_2_, 5% antimycotic/antibiotic, 1% Glutamax) for 2 h at 37°C and 5% CO_2_ to reduce bacterial contamination. Hookworms were then transferred to 2x substrate and incubated for a further 24 h at 37°C and 5% CO_2_ at a rate of ∼ 50 worms per 25 ml of media.

A total of 50 µg of *Ac*ESP from batch 1 was separated by SDS-PAGE and 18 bands were excised from the gel, reduced using dithiothreitol (DTT), alkylated with iodoacetamide (IAM) and digested with trypsin overnight as described previously [10]. One hundred (100) micrograms of *Ac*ESP from batch 2 was reduced and alkylated using DTT and IAM, respectively followed by trypsin digestion overnight. Peptides were separated using an OFFGEL fractionator as previously described [10]. All samples were desalted using C18 ZipTips after SDS-PAGE or Offgel separation. Peptides were analysed using a Shimadzu Prominance Nano HPLC coupled to an AB SCIEX Triple TOF+ 5600 mass spectrometer and processed using the software Analyst TF 1.6.1. The mass spectrometry proteomics data have been deposited to the ProteomeXchange Consortium via the PRIDE [11] partner repository with the dataset identifier PXD006511 and doi:10.6019/PXD006511.

Database searches were performed against a database consisting of the *A. caninum* genome and proteins from the common Repository of Adventitious Proteins (cRAP, http://www.thegpm.org/crap/) with Mascot using Mascot Daemon (v.2.5.1, Matrix Science) and X! Tandem (v.2015.12.15.2) and Comet (v.2016.01 rev.2) using PeptideShaker (v.1.11.0) [12].

A total of 237 and 289 proteins were identified with 2 or more peptides and FDR <1% using PeptideShaker and Mascot (Supplementary Table 1, 2), respectively, from which 211 were common between both search programs, while 26 and 78 were uniquely found by PeptideShaker and Mascot, respectively, resulting in a final quantity of 315 *Ac*ESP proteins (Supplementary Table 3). A total of 67 out of the 105 proteins identified by Mulvenna et al. [10] were also found in the present study (Figure 1A).

**Fig. 1.**
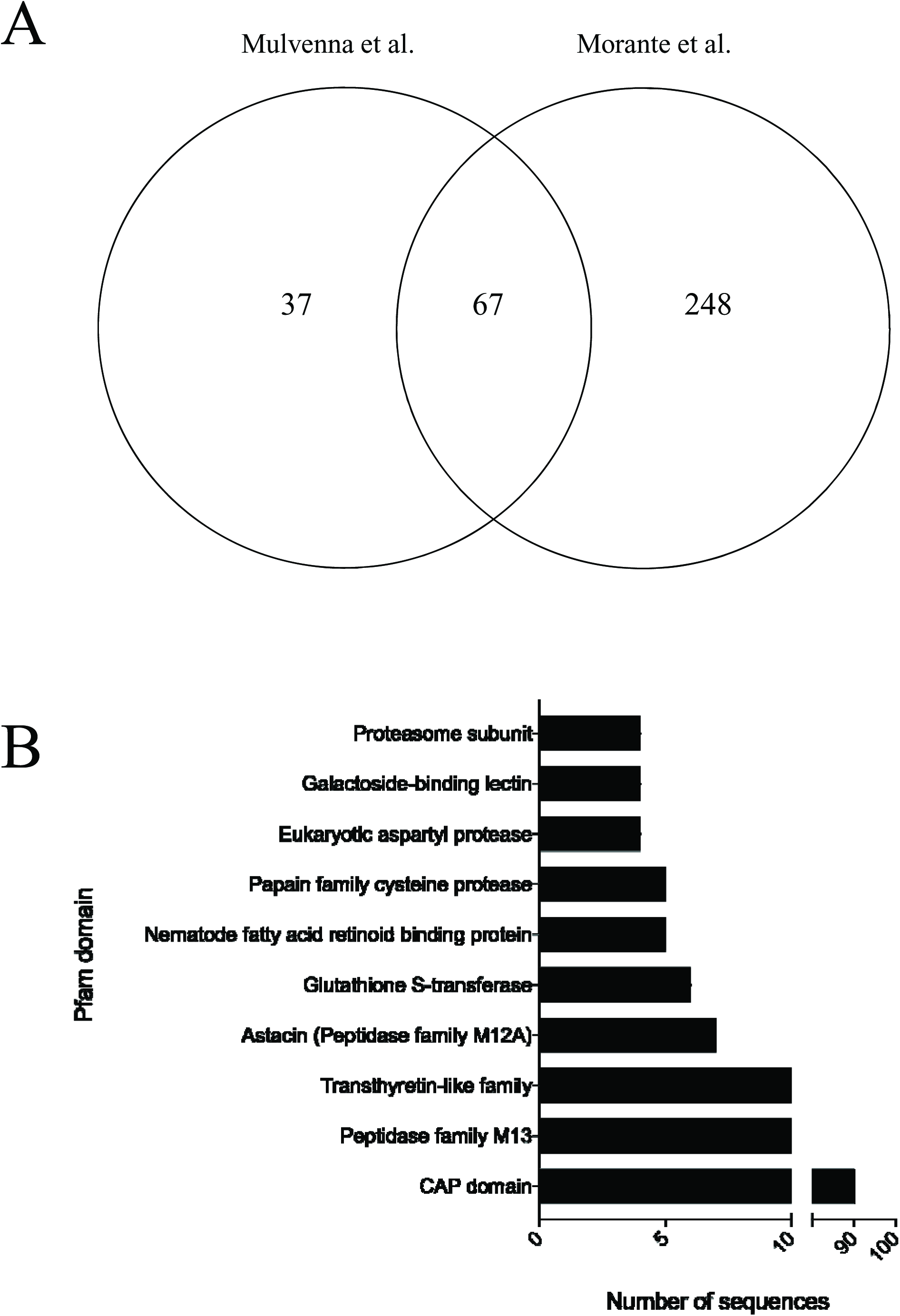
(A) Comparison between the number of *Ancylostoma caninum* excretory/secretory proteins (*Ac*ES) identified by Mulvenna et al. [10] and the present study. (B) Top ten most represented protein families in the *Ac*ES after a Pfam analysis. CAP: Cysteine-rich secretory protein family.

The top three most abundant proteins found by Mascot (Table 1) were ANCCAN_13497 (a previously described tissue inhibitor of metalloproteases; TIMP [13]), ANCCAN_25071 (a hypothetical protein), and ANCCAN_19759 (a sperm-coating protein; SCP, also called SCP/Tpx-1/Ag5/PR-1/Sc7 domain containing proteins; SCP/TAPS). The top three proteins found by X! Tandem and Comet using PeptideShaker (Table 2) were ANCCAN_03259 a platelet inhibitor, ANCCAN_01699 an SCP protein, and ANCCAN_13497 a TIMP (also found by Mascot in the top three most abundant proteins). TIMP proteins are a multifunctional family of inhibitors of matrix metalloproteases (MMP) associated with different functions in eukaryotic systems such as tissue remodelling, extracellular matrix turnover, cell proliferation and angiogenesis among others [14, 15]. However, in parasites, it has been previously suggested that TIMP-like proteins might not be functioning as MMP ihibitors [16]. The TIMP-like protein ANCCAN_13497 (previously identified as *Ac*-TMP-1 [13]) was already identified by Mulvenna et al. as the most abundant (and only TIMP-like) protein in the *Ac*ES [10]. Interestingly, other *A. caninum* TIMP-like protein, the renamed Anti-Inflammatory Protein-2 (AIP-2), has been shown to be a potent immunomodulatory protein that supresses airway inflammation in a mouse model of asthma [7]. AIP-2 has extensive homology with other TIMPs found in *Ac*ES, including the highly abundant ANCCAN_26655 and ANCCAN_24968, suggesting that these two proteins could have similar immunomodulatory properties. Four different TIMPs are present in *Ac*ES, and are highly abundant based on the emPAI and spectral count.

**Table 1.**
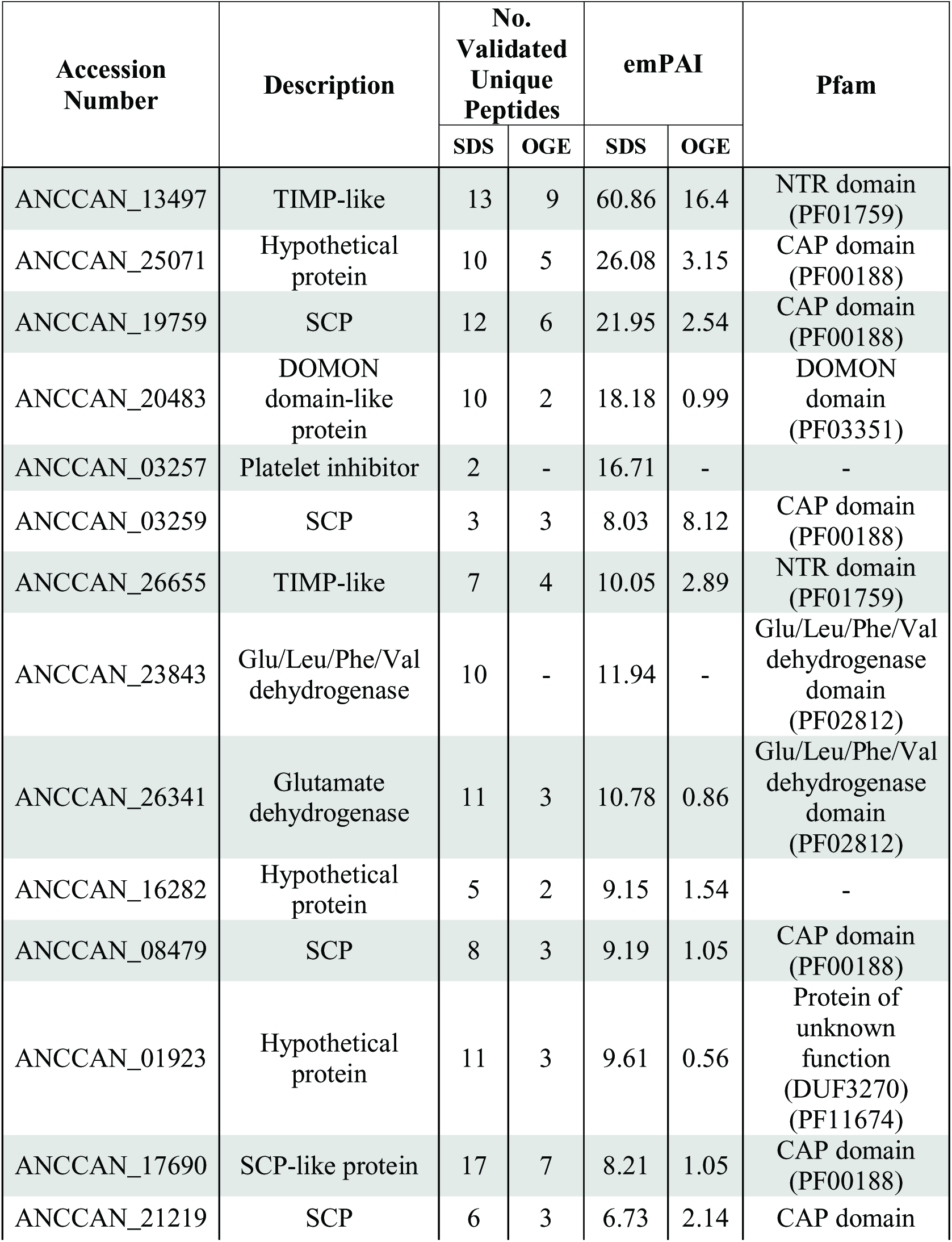
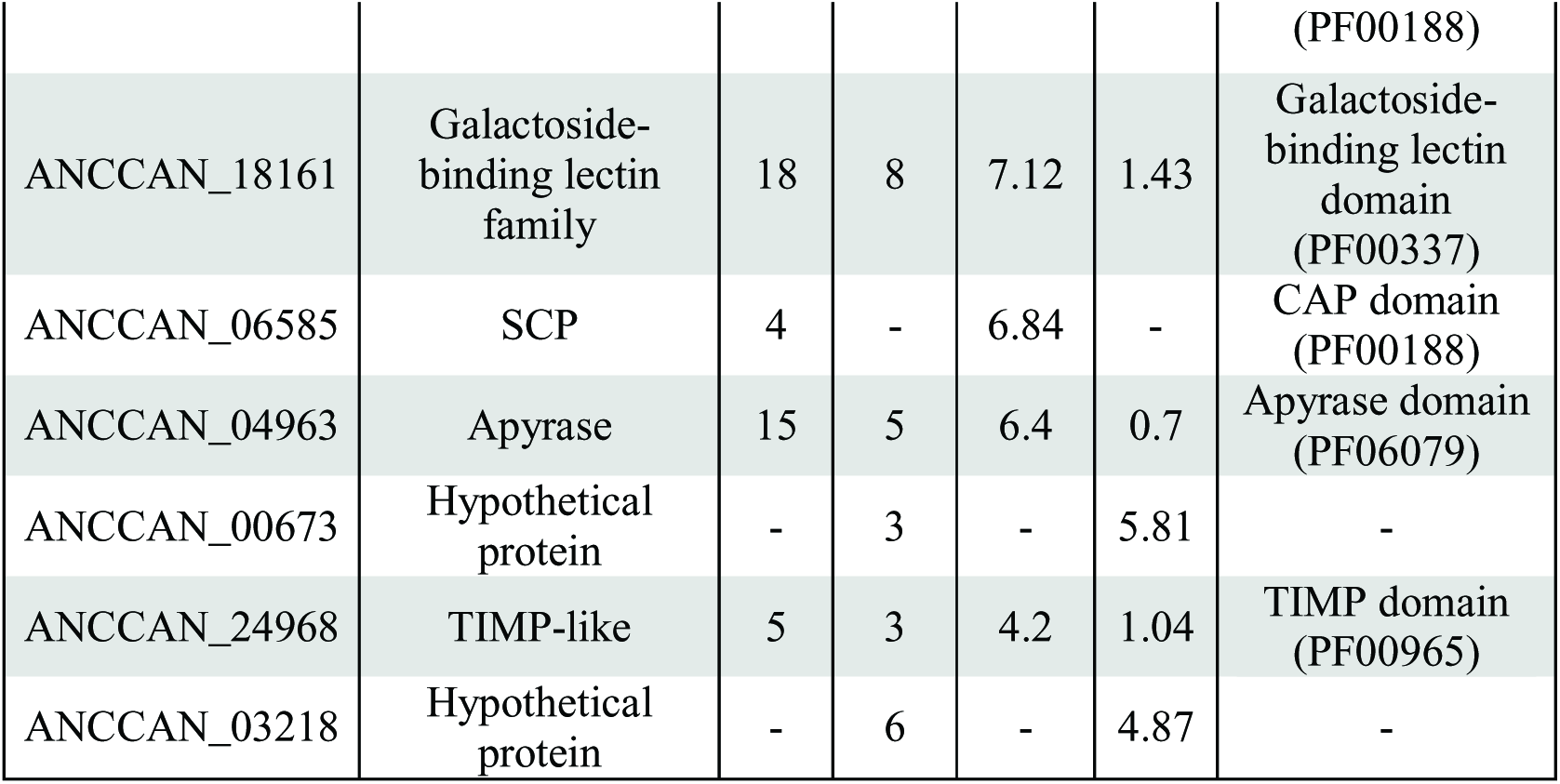
Top 20 proteins found by Mascot in the excretory/secretory products of *Ancylostoma caninum* adult worms based on emPAI. Proteins were identified by SDS-PAGE, Offgel, or both. CAP: Cysteine-rich secretory protein family; DOMON: dopamine beta-monooxygenase N-terminal; NTR: UNC-6/NTR/C345C module; SCP: sperm-coating protein; TIMP: tissue inhibitor of metalloproteases.

**Table 2.** Top 20 proteins found by X! Tandem and Comet in the excretory/secretory products of *Ancylostoma caninum* adult worms based on spectrum counting. Proteins were identified by SDS-PAGE, Offgel, or both. AnfO_nitrog: Iron only nitrogenase protein AnfO; CAP: Cysteine-rich secretory protein family; NTR: UNC-6/NTR/C345C module; SCP: sperm-coating protein; TIMP: tissue inhibitor of metalloproteases.

| Accession Number | Description | No. Validated Unique Peptides |  | Spectrum Count |  | Pfam |
| --- | --- | --- | --- | --- | --- | --- |
|  |  | SDS | OGE | SDS | OGE |  |
| ANCCAN_03257 | Platelet Inhibitor | 4 | 7 | 12605.4 | 3157.9 | - |
| ANCCAN_01699 | SCP | 3 | - | 9940.4 | - | CAP domain (PF00188) |
| ANCCAN_13497 | TIMP-like | 17 | 15 | 1674.2 | 4779.2 | NTR domain (PF01759) |
| ANCCAN_03254 | SCP-like protein | 7 | 5 | 2278 | 1622.4 | CAP domain (PF00188) |
| ANCCAN_03218 | Hypothetical protein | - | 5 | - | 2280.6 | - |
| ANCCAN_03214 | Hypothetical protein | 7 | - | - | 2244.7 | - |
| ANCCAN_23152 | Hypothetical protein | 3 | - | - | 2060.2 | AnfO_nitrog domain (PF09582) |
| ANCCAN_11008 | Secreted protein 4 precursor | 2 | 3 | 517.8 | 2046.8 | - |
| ANCCAN_20841 | Unknown | 4 | 3 | 495.5 | 1982.6 | - |
| ANCCAN_13591 | Hypothetical protein | - | 4 | - | 1869.4 | - |
| ANCCAN_16282 | Hypothetical protein | 5 | - | 1737.3 | - | - |
| ANCCAN_25071 | Hypothetical protein | 8 | 6 | 1395.2 | 1587.6 | CAP domain (PF00188) |
| ANCCAN_26655 | TIMP-like | 3 | 2 | 1567.9 | 1196.3 | NTR domain (PF01759) |
| ANCCAN_10664 | TIMP-like | - | 2 | - | 1455.6 | TIMP domain (PF00965) |
| ANCCAN_19423 | Hypothetical protein | - | 4 | - | 1238.5 | - |
| ANCCAN_05485 | SCP-like protein | 3 | 2 | 564.8 | 1093.2 | CAP domain (PF00188) |
| ANCCAN_18168 | Kunitz<br>Bovine<br>pancreatic<br>trypsin<br>inhibitor<br>domain | - | 3 | - | 1082.6 | Kunitz/Bovine<br>pancreatic<br>trypsin<br>domain<br>(PF00014) |
| ANCCAN_25718 | Excretory<br>secretory<br>protein 1 | 2 | 3 | 1060.7 | 1394.7 | - |
| ANCCAN_05778 | Copper zinc<br>superoxide<br>dismutase | 6 | 5 | 525.6 | 947.6 | Copper/zinc<br>superoxide<br>dismutase<br>domain<br>(PF00080) |
| ANCCAN_19037 | Hypothetical<br>protein | - | 4 | - | 881.3 | - |

The SCP proteins are highly represented in the infective larval stage of hookworms [17], where a role in larval penetration and infection has been hypothesized (reviewed by [18]). They have also been speculated to be involved in immunomodulation [19]. For instance, an *A. caninum* SCP-like protein (known as Neutrophil Inhibitory factor; NIF) is able to inhibit neutrophil function and oxidative stress [20]. Although we didn’t find this protein in the adult secretions, we have found two SCPs having extensive homology with the NIF sequence deposited in NCBI (accession number AAA27789.1): ANCCAN_04194 (88% identity and E-value = 6.9 ×ue^−89^) and ANCCAN_22933 (94.3% identity and E-value = 1.5 × 10^−65^). This could refer to a missannotation of the genome, since a blast search against the *A. caninum* genome using the deposited NIF sequence doesn’t return any sequence with 100% homology. SCP-related proteins (including SCPs and SCP-like proteins) contributed to 90 out of 315 (31%) of the overall protein families and were by far the most highly represented family of proteins in the *Ac*ES (Figure 1B). These results suggest that the SCP family might play a key role in orchestrating a parasitic existence and modulating the host’s immune response, and further studies should focus on this family of proteins.

Among the most represented protein domains in *Ac*ES was the metallopeptidase family M13 (11 proteins), the transthyretin-like family (10 proteins), astacin metalloproteases (7 proteins) and glutathione-s-transferases (GSTs; 6 proteins). The role of peptidases in the secretome of helminths have been linked to important roles in parasitism [21]; however, the roles of transthyretin-like proteins are still unknown (despite their abundance in nematodes in particular). Astacins are a family of metallopeptidases highly abundant in the secretome of helminths [22]. Indeed, the human hookworm *N. americanus* has 82 astacin-encoding genes [23], and astacins are abundantly represented in ESPs of the rat hookworm *Nippostrongylus brasiliensis* [24] and other nematodes [25]. The role of astacins is not fully understood, although, in nematodes, they have been shown to participate in host invasion and parasite development [26, 27]. It is also noteworthy to highlight the presence of GSTs in the *Ac*ES proteome. *Na*-GST-1, a GST secreted by adult stage *N. americanus* that is thought to play a role in feeding by detoxifying the free heme produced after hemoglobin ingestion, is being currently tested as a vaccine against *N. americanus* [28]. Other molecules that have been tested as vaccines against hookworms include aspartic proteases [29] and cysteine proteases [30], which are protein families also represented in our study (Figure 1B). In the present study we found 5 different cysteine proteases (ANCCAN_06649, ANCCAN_06644, ANCCAN_30567, ANCCAN_06619 and ANCCAN_06647) and four aspartic proteases (ANCCAN_18339, ANCCAN_13546, ANCCAN_29459 and ANCCAN_25067) Thus, it is tempting to speculate that the GSTs, cysteine proteases and aspartic proteases identified in the present study could be potential vaccines against the dog hookworm.

In the present study we have reanalysed the protein constituents of *Ac*ES in order to gain a more comprehensive snapshot of the hookworm secretome and how this impacts on host-pathogen interactions. We have identified almost three times as many proteins as previously reported in the ESP of this important parasite. In addition, new proteins of interest with potential as both novel immunoregulatory biologics and vaccine candidates have been identified, and clearly warrant future exploration.

## ACKNOWLEDGMENTS

This work was supported by a program grant from the National Health and Medical Research Council (NHMRC) [program grant number 1037304] and a Senior Principal Research fellowship from NHMRC to AL (1117504). The funders had no role in study design, data collection and analysis, decision to publish, or preparation of the manuscript. The authors declare no competing financial interests.

